# Diffusion model-based image generation from rat brain activity

**DOI:** 10.1101/2024.05.26.595934

**Authors:** Kotaro Yamashiro, Nobuyoshi Matsumoto, Yuji Ikegaya

## Abstract

Brain-computer interface (BCI) technology has gained recognition in various fields, including clinical applications, assistive technology, and human-computer interaction research. BCI enables communication, control, and monitoring of the affective/cognitive states of users. Recently, BCI has also found applications in the artistic field, enabling real-time art composition using brain activity signals, and engaging performers, spectators, or an entire audience with brain activity-based artistic environments. Existing techniques use specific features of brain activity, such as the P300 wave and SSVEPs, to control drawing tools, rather than directly reflecting brain activity in the output image. In this study, we present a novel approach that uses a latent diffusion model, a type of deep neural network, to generate images directly from continuous brain activity. We demonstrate this technology using local field potentials from the neocortex of freely moving rats. This system continuously converted the recorded brain activity into images. Our end-to-end method for generating images from brain activity opens up new possibilities for creative expression and experimentation.

## 1 Introduction

Brain-computer interface (BCI) technology has witnessed remarkable advancements in recent years, revolutionizing various domains such as clinical applications, assistive technology, and human-computer interaction research (1–5). By establishing a direct communication pathway between the human brain and external devices, BCI systems enable users to control and interact with their surroundings using neural activity signals. BCI has shown great potential in motor rehabilitation by enabling individuals with motor disabilities to control external devices using their brain signals (2,6–8). For example, stroke patients with impaired hand function can use BCI to control a robotic exoskeleton or a virtual avatar, allowing them to perform rehabilitative exercises and regain motor skills (9–12). BCI have also made advancement in assistive communication, where individuals with severe motor disabilities, such as those with amyotrophic lateral sclerosis (ALS) or locked-in syndrome can communicate by translating their brain activity into text or speech output (13–15). By focusing on specific tasks or commands, users can spell out words, select options on a screen, or even generate synthetic speech. BCI has also shown promise in the field of mental health by providing insight into brain activity patterns associated with various conditions such as depression, anxiety, and attention deficit hyperactivity disorder (ADHD) (16–24). Neurofeedback using BCI allows individuals to self-regulate their brain activity by providing real-time feedback. This technique can help train individuals to increase or decrease specific brain wave patterns, leading to improved cognitive function and emotional well-being (3).

Notably, the potential of BCI technology extends beyond functional applications, as it has also gained traction in applications for healthy individuals. Some of these applications include controlling smartphone apps with EEG and playing games using brain activity. BCI has also created a link between brain activity and art (16,25). Recently, artists have used BCI to create art that is dynamically influenced by their own brain activity or that of the audience. Some of this art includes exhibitions that are influenced by the audience’s brain activity and music that is created according to the measured activity (26–29). The process usually involves an intermediate step that decodes the user’s intentions and then adjusts the palette based on the decoded actions. While this painting BCI provides users with a novel way to create art, the experience can be enhanced by allowing users to seamlessly transform their brain activity into paintings in a single step, free from intentional intervention. Painting with decoded intent will limit the user’s creations to what they can consciously imagine. By completely removing painting output from conscious control, entirely new forms of art can emerge.

In recent years, there has been a growing interest in diffusion models (DMs), which are deep generative models (30–32). The DMs have shown remarkable performance in various tasks, such as text conditioned image generation (33–35), upsampling image resolution (36,37), and image colorization (38,39). To make training and inference more efficient, latent diffusion models (LDMs) have been introduced, which compute diffusion processes in the latent space generated by autoencoders (40). This results in reduced computational costs and enables the generation of high-resolution images. Diffusion models have expanded the boundaries of art by introducing new ways to create and manipulate visual content.

In this paper, we present a novel approach that leverages LDMs to generate images directly from brain activity signals. Our method provides an end-to-end solution that seamlessly transforms neural activity into visual art, eliminating the need for intermediate steps or manual interpretation. By presenting our experimental results with recorded brain activity from freely moving rats, we demonstrate the feasibility and potential of our approach for translating neural signals into visual artistic output.

## 2 Related Work

The exploration of creative expression through the fusion of art and neural processes has been the subject of research (41,42), including modalities such as music and painting. Overall, this translating framework first extracts the subject’s intentions, which are then used to control various interfaces, such as an audio mixer and a painting palette (26–28,43). Previous research has attempted to help people with disabilities and has successfully enabled paralyzed patients to express their artistic creativity.

### 2.1 Composing Music with Neural Signals

Musical composition is one of the transformations of neural activity into a form of art (25,44). In the context of these investigations, the transformation of neural signals into music is based on the analysis of electroencephalography (EEG) data. Specifically, previous studies have focused on EEG power spectrums, as well as the topological patterns exhibited by EEG data over an array of electrode assemblies. This computed power spectrum is used to control audio mixing and the tuning of musical parameters (43). It can also facilitate direct engagement with musical instruments, thereby enriching the interaction between individuals and their musical creations.

### 2.2 Painting using Neural Signals

Painting using neural activity is another application that enables creative expression. In these paradigms, a specific neural component triggered by an internal or external event is used. The conversion of neural activity into painting is accomplished by decoding of the subject’s intentions from the specific neural component. The subject’s intentions are then used to control a palette on a screen. In previous research, two types of neural components have been considered, the P300 wave (26–28) and the steady state visual evoked potential (SSVEP) (29).

The P300 wave is detectable as a positive shift in the EEG signal 200-400 ms after the external stimulus is presented to the subject. The P300 wave is associated with the subject’s response to a stimulus and is considered an endogenous potential. The application of P300 waves to BCI is most commonly used as a speller for locked-in patients. The P300 speller uses a speller matrix containing letters and digits. In this system, the user directs attention to a character within a matrix, where each row or column is rapidly highlighted in a pseudo-random manner. If the highlighted row or column contains a letter desired by the subject, the P300 wave is detected in the EEG signal. By analyzing the evoked responses, the BCI is able to identify the desired letter by determining the row and column that evoke the most significant response. P300 brain painting uses this P300 speller matrix to create the P300 palette matrix, where different colors and shapes can be selected from the P300 response. The drawback of the P300 painting application was the slow response, which affected the efficiency of the system.

Some researchers have turned to SSVEP-based applications to overcome the challenges of P300 systems. The SSVEP is a continuous response generated in the visual cortex that synchronizes with the frequency of visual stimuli presented to the subject. The subject is exposed to a matrix of cells, each flickering at a different frequency. By directing attention to a specific target cell, the subject can effectively make a choice. This SSVEP-driven approach provides an enhanced rate of information transfer. The integration of hybrid brain-computer interface (BCI) systems that combine P300 and SSVEP has led to advances in real-time human-computer interaction. The brain painting application that used this system has successfully improved the subject’s real-time interaction.

Previous research on brain painting systems has effectively enabled users to express their artistic creativity by facilitating the translation of their intentions into artistic compositions. These systems typically begin by extracting the subject’s intentions, which are then used as input to control the painting palette on a computer screen. Consequently, the resulting artwork represents the subject’s intentions. Thus, the development of a novel application of brain painting depends on the development of a system that directly translates neural signals into works of art, which represents a promising avenue for artistic expression in BCI.

## 3 Methods

### 3.1 Datasets

#### 3.1.1 Ethical approvals

Animal experiments were performed with the approval of the Animal Experiment Ethics Committee at the University of Tokyo (approval number: P29-7) and according to the University of Tokyo guidelines for the care and use of laboratory animals. These experimental protocols were carried out in accordance with the Fundamental Guidelines for the Proper Conduct of Animal Experiments and Related Activities of the Academic Research Institutions (Ministry of Education, Culture, Sports, Science and Technology, Notice No. 71 of 2006), the Standards for Breeding and Housing of and Pain Alleviation for Experimental Animals (Ministry of the Environment, Notice No. 88 of 2006) and the Guidelines on the Method of Animal Disposal (Prime Minister’s Office, Notice No. 40 of 1995). Although our experimental protocols require that animals be humanely euthanized if they show signs of pain, marked lethargy, and discomfort, we did not observe such symptoms in any of the rats tested in this study. Every effort was made to minimize animal suffering.

#### 3.1.2 Animal Strains

Local field potentials (LFPs) were recorded from Long-Evans rats (Japan SLC, Shizuoka, Japan). They were individually housed under conditions of controlled temperature and humidity (22 ± 1°C, 55 ± 5%) and maintained on a 12:12-h light/dark cycle (lights off from 7:00 to 19:00) with *ad libitum* access to food and water unless otherwise specified.

#### 3.1.3 Surgery

A custom 32-channel electrode was used to record LFPs from rats. This electrode was designed to cover the right S1 region, specifically the forelimb and hindlimb subregions. At the beginning of surgery, a rat was anesthetized with 2-3% isoflurane gas. A square craniotomy (2-6 mm posterior and 1-5 mm lateral to bregma) was performed using a dental drill. In addition, two stainless steel screws were implanted in the bone above the cerebellum as ground and reference electrodes. The recording device and electrodes were secured to the skull with stainless steel screws and dental cement. After surgery, the rat was housed in a transparent Plexiglas cage with free access to food and water for one week.

#### 3.1.4 Recording

After one week of recovery from surgery, LFPs were recorded from a freely moving rat in an open field. The rat was placed in a 40×40 cm open field. Several plastic objects were placed inside the open field to motivate the rat to move and not stay in one place. LFPs were recorded using the Open Ephys recording system (http://open-ephys.org). The recorded LFPs were referenced to the ground and digitized at 3000 Hz. The digitized data were simultaneously recorded with an overhead camera at 30 Hz.

### 3.2 Latent Diffusion Models

Diffusion models (DMs) are probabilistic generative models trained to recover a sample variable from Gaussian noise. Given a Gaussian noise input, the model transforms it into a sample from a learned data distribution through an iterative denoising process. The diffusion model is defined by two processes, the forward diffusion process and the backward diffusion process. To further explain this concept, we will review the mechanics of the diffusion process.

The forward diffusion process starts with a sample from the known data distribution. Small amounts of noise, usually Gaussian noise, are added to this initial sample stepwise. The addition of noise at each step produces a sequence of noisy samples. For each step *t* of the original data, Gaussian noise is added incrementally up to step *T*. The larger the *T*, the final *x*_*T*_ will be nearly an isotropic Gaussian distribution.

In the backward diffusion process, the diffusion model learns the reverse process of the forward diffusion process to acquire a sample from the Gaussian distribution. Essentially, at each step in this backward diffusion process, a noisy sample *x*_*T*−1_ is predicted based on the preceding sample *x*_*T*_. This recovery of the original data from the Gaussian distribution is accomplished through the implementation of a neural network. The widely adopted architecture for this neural network in the reverse diffusion process is the U-net. Once the model is trained to recover data from the Gaussian distribution, the model is capable of generating a new sample from the original data distribution given a noisy input. The application of diffusion models to images has allowed the generation of realistic images from initial Gaussian noise data.

A variant of diffusion models known as LDM is used in this research. In conventional diffusion models, image generation takes place in high-dimensional pixel space, which requires significant computational resources. In contrast, the LDM performs diffusion processes within the latent diffusion space, a compressed image representation generated by an autoencoder. Initially, Gaussian noise is compressed into this latent space by an encoder, preserving maximum information while reducing sample dimensions. The subsequent diffusion process mirrors that of standard diffusion models. Finally, a decoder reconstructs an output image based on a sample from the latent space.

In this study, the LDM used is known as OpenJourney, a fine-tuned version of Stable Diffusion version 1.5 trained on a large image dataset. The original model was adapted to manipulate generated images based on textual input via a pre-trained text encoder (Contrastive Language-Image Pretraining: CLIP). However, in our implementation, no conditioning by textual input was applied. Consequently, the generated images were entirely dependent on the input neural activity.

### 3.3 Seamless Transformation of Neural Signals to Images

To convert the LFP into images, the LFP was segmented and processed through a denoising U-net and a latent-to-image decoder provided by the author of the LDM (Fig. 1A). The LDMs require input data with dimensions of 64×64×4. First, rat LFPs were filtered with a low-pass filter at 30 Hz. The filtered signal was segmented into windows of 512 points with a sliding step of 100, creating a continuous data matrix of 512×32 points every 1/30 second (Fig. 1B). This matrix was then resized to 64×64×4 and shuffled with a fixed random seed, mapping the segmented LFP to a noisy latent vector zt. The fixed seed ensured that changes in the same pixel in successive data matrices were aligned with the temporal signal of the LFP. The noisy latent vector zt was processed by denoising U-Net to produce a denoised latent vector z. In this process, text input is commonly inserted into the denoising process for text conditioning. However, our implementation deliberately excluded any text input, thus generating images that only reflect the fluctuation of LFPs. Finally, the denoised latent vector was converted to image space using the latent-to-image decoder. The resulting image sequence was transformed into a 30 fps video clip. The model was executed using a Python script adapted from the official StableDiffusion GitHub repository and ran on the NVIDIA RTX A6000 platform. Default parameters provided by the authors of LDM were used in all implementations unless otherwise noted.

**Figure 1.**
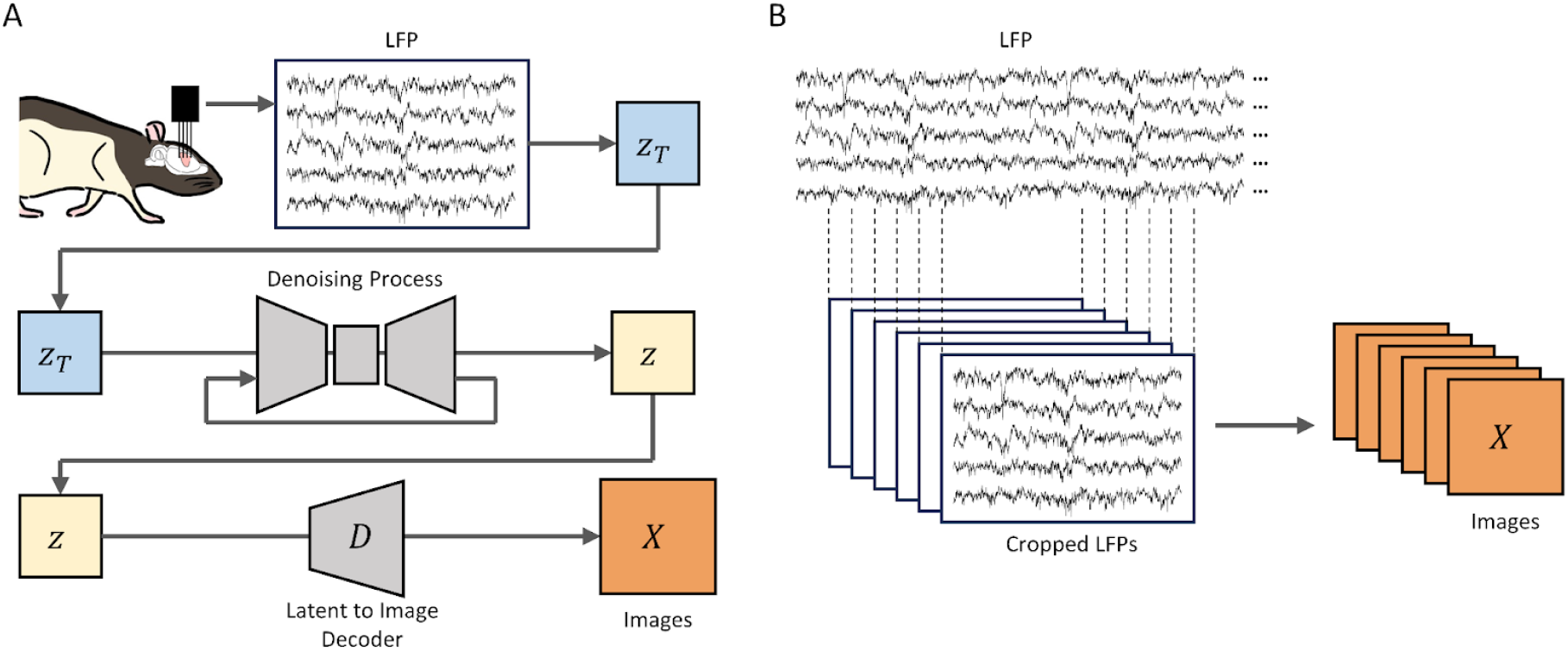
Overview of the method used to continuously generate images from rat LFP. **A**. Detailed illustration of the steps taken to generate image from rat LFP. The recorded LFPs were mapped onto noisy latent vector ***z***_***T***_, which was then processed by denoising U-Net to produce a denoised latent vector ***z***. This latent representation of the denoised image was then processed by a latent to image decoder to produce image ***X*. B**. Schematic of the generation of morphing images from rat LFP. A 512×32 matrix was sampled every 1/30 second with overlaps. The cropped LFPs were then processed through the denoising U-Net and the decoder to generate images at a rate of 30 Hz.

## 4 Results and Discussion

In this work, we developed a pipeline that enables the generation of art from brain activity using LDM (Fig. 1A). In this pipeline, the recorded brain activity (LFPs, herein) was mapped to a latent vector, which was then processed by a denoising U-net. The denoised latent vector was then expanded into image space by a latent-to-image decoder. This entire process runs seamlessly without any user or operator input or intervention, enabling a fully automated generation of images derived from brain activity.

We have further extended our LFP-to-image transformation framework to account for the continuous fluctuation of brain activity. Fluctuations in brain activity are experimentally observed and can be characterized as non-deterministic and deterministic, along with oscillations at different frequencies (45–48). In the cortex, these stochastic processes are known to be non-Gaussian (49) and unpredictable. We applied our LFP-to-image transformation to capture the fluctuations in continuously changing LFPs to visualize the stochastic process in the rat cortex (Fig. 1B). In this approach, we first extracted LFP segments from the continuous recording with a sliding step of 100 points, resulting in similar matrices when they were close together. Consequently, each matrix was positioned close to its neighboring matrices in multidimensional space. To illustrate the arrangement of these matrices, we used Uniform Manifold Approximation and Projection (UMAP) on the converted LFP matrix (Fig. 2A). Since the noisy latent vectors converted from segmented LFPs also reflected continuity in multidimensional space, the output of the LDM was also similar between successive frames. This allowed the generation of morphing images that captured fluctuations in brain activity in the rat cortex (Fig. 2B).

**Figure 2.**
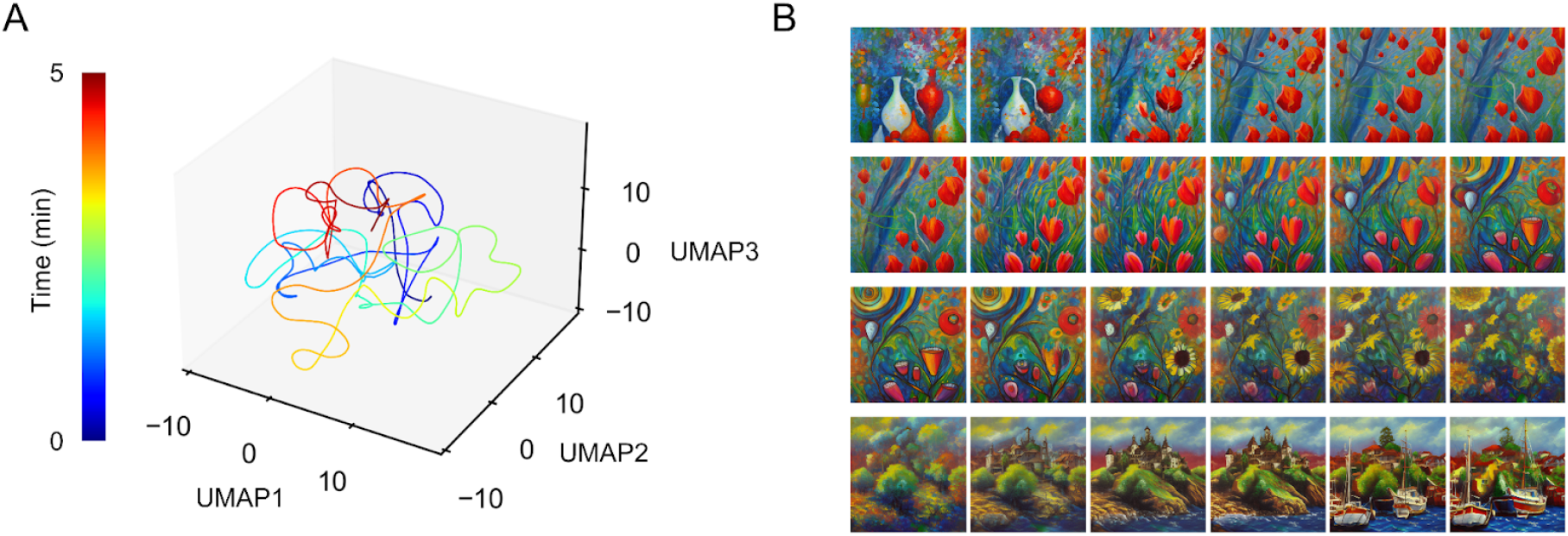
Examples of morphing images generated from rat LFPs. **A**. 3D visualization of latent vector ***z***_***T***_ using UMAP. Embeddings are from a 5-minute recording of LFPs. Pseudocolor labels indicate the elapsed time since the start of the recording. **B**. Samples from morphing images over a 10-s duration.

Because the global state of neural activity is constantly changing within the multidimensional neural manifold(50,51) and combined with local dynamics such as oscillation strength and subtle shifts in neuronal firing patterns (52,53), similar images are rarely generated, especially when their latent vectors are temporally separated. Consequently, our LFP-to-image pipeline has the potential to generate an infinite number of image variations (Fig. 3). Furthermore, because open-source models can be downloaded and fine-tuned for LDM customization, a wide variety of LDMs are readily available today. As an example of the customization of our model, Figure 4 shows an example of changing the style of the generated images. While our primary focus has been on image generation using LDMs, recent advances have broadened the scope to include the generation of audio through diffusion processes (54,55). It is conceivable that audio-generating LDMs could be integrated with brain activity recordings, allowing the composition of music that reflects fluctuations in neural activity. This wealth of possibilities underscores the virtually limitless potential for art generation using BCI technology.

**Figure 3.**
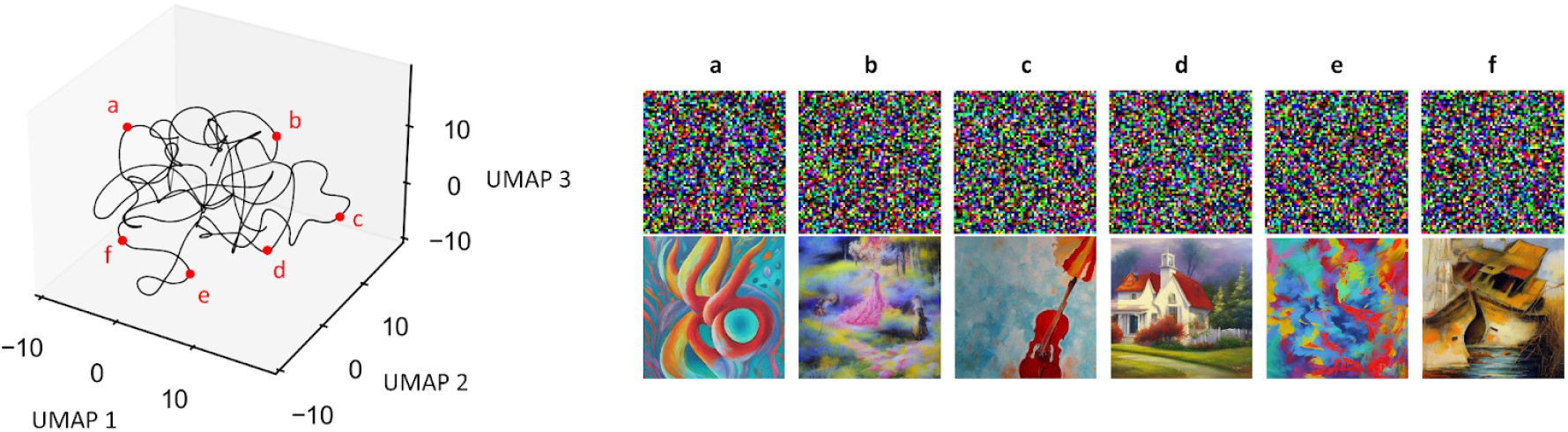
Temporally spaced sample selection from a 5 min LFP recording. ***Left***: Spatial distribution of selected frames in the embedding space. ***Right***: Noisy latent vector ***z***_***T***_ and the corresponding generated image obtained by the diffusion process and the latent-to-image decoder.

**Figure 4.**
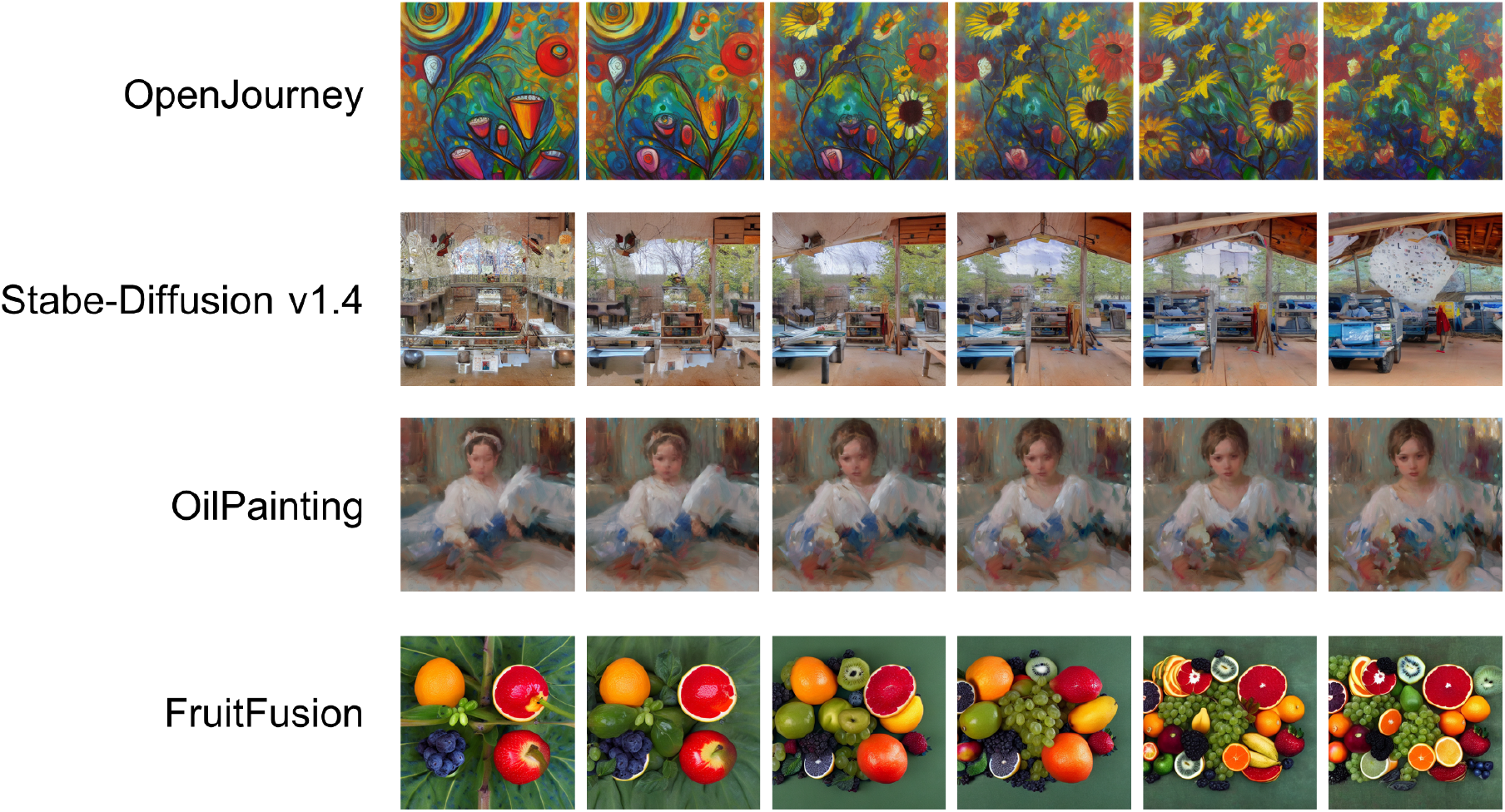
Various styles of generated images. Morphing images generated from LFPs using different models. From top to bottom: OpenJourney, Stable-Diffusion v1.4, OilPainting, FruitFusion. All are available on Civitai.com.

If our framework were extended to humans, the potential to generate works of art without the need for conscious control holds promise for enhancing creativity (56,57). The integration of art and BCIs has advanced in recent years, spanning domains such as music (44,58–61) and painting (56,57,62). In brain painting, previous research has successfully used characteristic responses in brain activity to directly control a palette for drawing paintings (27–29,57,63,64). Our approach differs from other work on image generation in that it eliminates the need for an intermediate step in decoding intentions. The seamless transformation of brain activity into images provides a unique experience for the audience. In addition, our system may make it easier for artists to generate creative ideas because the output is independent of their consciousness. Physiological measures of creative processes are often assessed using divergent thinking (65) and remote association (66,67). The unconscious generation of images from one’s own brain activity introduces a novel concept to the user. For example, divergent thinking is a cognitive process used to generate creative ideas by exploring numerous potential solutions. Offering a new perspective on painting through the capabilities of artificial neural networks expands the range of solutions beyond conscious ideas, thereby fostering heightened creativity.

The fusion of art and BCI technology offers innovative possibilities, as evidenced by our development of frameworks that enable the direct translation of brain activity into visual art. Although our implementation has been in rats, the concept of “brain painting” not only provides a novel means of expression for artists and non-artists alike, but also offers a unique avenue for the unconscious generation of creative ideas. At the intersection of neuroscience, artificial intelligence, and artistic creation, the continuing evolution of technology promises not only to redefine our approach to art, but also to unlock new dimensions of creativity.

## Acknowledgments

The authors thank the laboratory members for helpful discussions, valuable comments, and technical support.

